# Queuosine Salvage in *Bartonella henselae* Houston 1: A Unique Evolutionary Path

**DOI:** 10.1101/2023.12.05.570228

**Authors:** Samia Quaiyum, Yifeng Yuan, Guangxin Sun, R. M. Madhushi N. Ratnayake, Geoffrey Hutinet, Peter C. Dedon, Michael F. Minnick, Valérie de Crécy-Lagard

## Abstract

Queuosine (Q) stands out as the sole tRNA modification that can be synthesized via salvage pathways. Comparative genomic analyses identified specific bacteria that showed a discrepancy between the projected Q salvage route and the predicted substrate specificities of the two identified salvage proteins: 1) the distinctive enzyme tRNA guanine-34 transglycosylase (bacterial TGT, or bTGT), responsible for inserting precursor bases into target tRNAs; and 2) Queuosine Precursor Transporter (QPTR), a transporter protein that imports Q precursors. Organisms like the facultative intracellular pathogen *Bartonella henselae*, which possess only bTGT and QPTR but lack predicted enzymes for converting preQ_1_ to Q, would be expected to salvage the queuine (q) base, mirroring the scenario for the obligate intracellular pathogen *Chlamydia trachomatis*. However, sequence analyses indicate that the substrate-specificity residues of their bTGTs resemble those of enzymes inserting preQ_1_ rather than q. Intriguingly, mass spectrometry analyses of tRNA modification profiles in *B. henselae* reveal trace amounts of preQ_1_, previously not observed in a natural context. Complementation analysis demonstrates that *B. henselae* bTGT and QPTR not only utilize preQ_1_, akin to their *Escherichia coli* counterparts, but can also process q when provided at elevated concentrations. The experimental and phylogenomic analyses suggest that the Q pathway in *B. henselae* could represent an evolutionary transition among intracellular pathogens—from ancestors that synthesized Q *de novo* to a state prioritizing the salvage of q. Another possibility that will require further investigations is that the insertion of preQ_1_ has fitness advantages when *B. henselae* is growing outside a mammalian host.

**Author summary:** Transfer RNAs (tRNAs) are adaptors that deliver amino acids to ribosomes during translation of messenger RNAs (mRNAs) into proteins. tRNA molecules contain specially-modified nucleotides that affect many aspects of translation, including regulation of translational efficiency, as modified nucleotides primarily occur near the portion of tRNA (anticodon) that directly interacts with the coding sequence (codon) of the mRNA while it is associated with a ribosome. Queuosine (Q) is a modified tRNA nucleotide located in the anticodon that can be synthesized or uniquely imported from the environment as Q or a precursor using a salvage mechanism. Free-living bacteria, e.g., *E. coli*, can synthesize Q or salvage precursors from the environment, but many obligate intracellular pathogens, e.g., *Chlamydia trachomatis*, cannot synthesize Q and must import a precursor from eukaryotic hosts. In this study, we determined that *Bartonella henselae*, a facultative intracellular bacterial pathogen of vascular cells, falls somewhere in the middle, as it is unable to synthesize Q but can salvage Q or certain precursors. The unusual nature of *Bartonella*’s system suggests different evolutionary scenarios. It could be a snapshot of the transition from Q synthesis to strict Q salvage or represent a unique adaptation to a complex multi-host lifestyle.

## Introduction

tRNA modifications fine-tune translation by various mechanisms such as modulating the efficiency or accuracy of translation or affecting tRNA stability (1). Recent studies have revealed that modifications can play key roles in bacterial pathogenesis (2,3). Queuosine (Q) is a modification of the wobble base in tRNAs with GUN anticodons of many bacteria and eukaryotes that can affect both translational efficiency and accuracy depending on the organism (4,5). The *in vivo* significance of this modification has remained enigmatic for decades as it has been lost repeatedly during evolution (6), but recent studies have suggested that it may act as a regulatory component in the translation of proteins derived from genes enriched in TAT codons compared with TAC codons (7,8). In bacteria, Q modification was shown to have roles in oxidative stress, metal homeostasis, and virulence (9–12).

Q is synthesized from guanosine triphosphate (GTP) by bacteria in a complex eight-step pathway fully elucidated in *E. coli* (Fig. 1A). Four enzymes (GCHI, QueD, QueE, and QueC) convert GTP into 7-cyano-7-deazaguanine (preQ_0_). QueF then reduces preQ_0_ to 7-aminomethyl-7-deazaguanine (preQ_1_) that is inserted into tRNAs by tRNA guanine-34 transglycosylase (bacterial TGT, or bTGT) (4). The inserted base preQ_1_ is converted to Q by two additional steps involving QueA and QueG or QueH, depending on the organism (4,13). It should be noted that Q is the only tRNA modification that can be salvaged or recycled (5). Eukaryotes only use the salvage route, and their TGT enzyme (eTGT), a heterodimeric QTRT1/QTRT2 complex, incorporates the queuine (q) base directly into the target tRNAs (5). The salvage routes in bacteria vary greatly. Some organisms lack the preQ_0_ or preQ_1_ pathway genes but encode all the downstream genes and import these precursor bases to finalize Q synthesis *in vivo* (14). This salvage route is also observed in organisms like *E. coli* that can synthesize Q *de novo*. Other bacteria lack all canonical synthesis genes except *tgt* and can salvage q like eukaryotes. Two q salvage routes have been identified in bacteria, to date (15), including the direct salvage route found in the intracellular pathogen *Chlamydia trachomatis*, or the indirect salvage route found in the gut microbe *Clostridioides difficile* (Fig. 1B and C, respectively). In the direct route, the substrate specificity of the *C. trachomatis* bTGT enzyme has shifted to insert q instead of preQ_1_ like most bacterial homologs (15). In the indirect route, a recently discovered radical enzyme (QueL) can regenerate the preQ_1_ intermediate from a q precursor that is imported directly or derived from the hydrolysis of the Q nucleoside by QueK (15). Only a few transporters involved in Q salvage pathways have been identified and experimentally characterized. The first was the YhhQ /COG1738 family, now renamed QPTR (Queuosine Precursor Transporter), involved in the transport of preQ_0_ and preQ_1_ in *E. coli* and q in *C. trachomatis* (14,15). Members of the Energy-Coupling Factors (ECFs) family had been predicted to be involved in preQ_0_ and preQ_1_ transports (15)) and two of the *C. difficile* specificity components (or ECF-QueT) were shown to transport a variety of Q precursors in a reconstituted system (15).

**Fig 1.**
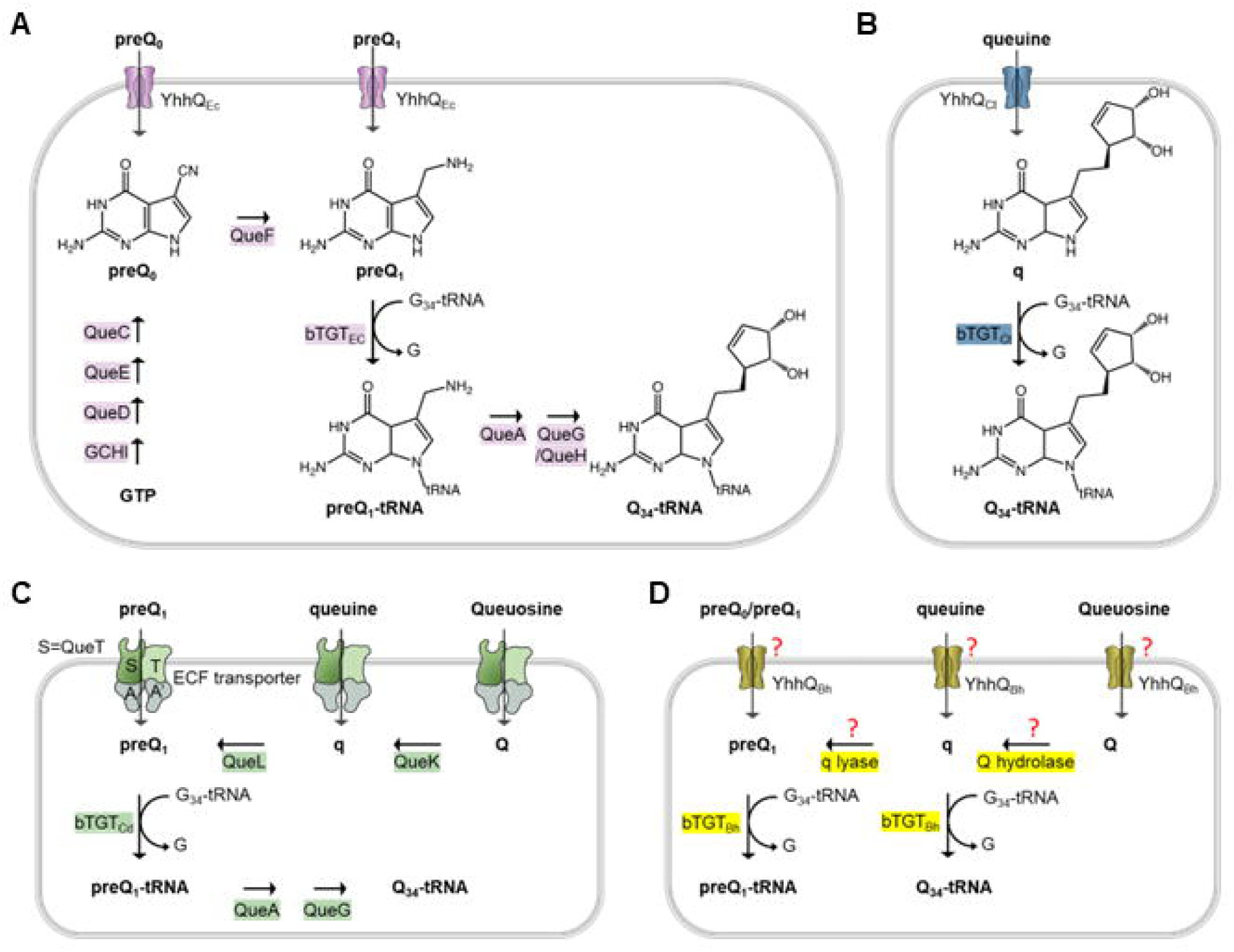
Known or predicted Q synthesis pathways. (A) preQ_0_/preQ_1_ synthesis and salvage pathways in *E. coli*; (B) q salvage pathway in *C. trachomatis*; (C) preQ_1,_ q and Q salvage pathways in *C. difficile*. The ECF transporters include 4 subunits: S, the substrate-specific transmembrane component (QueT); T, the energy-coupling module; A and A’, a pair of ABC ATPase. (D) Possible q and Q salvage pathways in *B. henselae* Houston 1.

The proportion of Q-modified tRNAs can change with the developmental stage in several eukaryotic parasites that undergo complex cycles that switch between hosts such as *Trypanosoma cruzei* and *Entamoeba histolytica* (16,17). Very little is known about the role of Q in bacterial pathogens that also switch between mammalian and insect hosts such as the facultative intracellular pathogen *Bartonella henselae*. This bacterium uses fleas and possibly ticks as vectors during blood feeding. Feces of these insects can also infect cats when they are scratched into a break in the skin. Once inside the cat, the bacteria enter the bloodstream, primarily residing within endothelial cells, where they multiply (18). *B. henselae* exhibits a high level of heterogeneity (19) and 16S rRNA sequence analyses led to the identification of two serotypes, Houston-1, and Marseille (20,21). Metabolic reconstruction of the Q synthesis pathway of *B. henselae* Houston 1 suggested it utilized a direct q salvage pathway like *C. trachomatis*. However, analysis of *B. henselae* bTGT and QTPR sequences did not match this prediction as these were more similar to the bTGT and QPTR enzymes that recognize preQ_1_. To resolve this discrepancy, we set out to characterize experimentally the Q salvage enzymes in this facultative intracellular pathogen.

## Results

### Metabolic reconstruction and sequence analyses of queuosine salvage genes give contradictory results

We previously showed that QPTR proteins have different substrate specificities, shifting from preQ_0_ and preQ_1_ in *E. coli* to q in *C. trachomatis* (14,15). To better understand the molecular determinants that drive this change in specificity, we constructed a Sequence Similarity Network (SSN) of the QPTR family (PF02592). We then colored the SSN based on the presence/absence of the Q synthesis genes in the corresponding genome as an indirect way to predict the substrate specificity of a given QPTR protein. As shown in Fig. 2, we were able to generate an SSN that separated the QPTR’s predicted to salvage preQ_0_ and preQ_1_ (in yellow or red) from those predicted to salvage q (in blue). However, a few exceptions stood out even at a stringent alignment score of 70. The QPTRs found in organisms that encode only *tgt*, and hence predicted to salvage q, cluster with QPTR proteins predicted to salvage preQ_0_/ preQ_1_ because their genomes encode QueA and QueG or QueH (circled in Fig. 2). These include QPTR proteins from *Bartonella* species such as *B. henselae* Houston-1 (UniProt id A0A0H3M726_BARHE).

**Figure 2.**
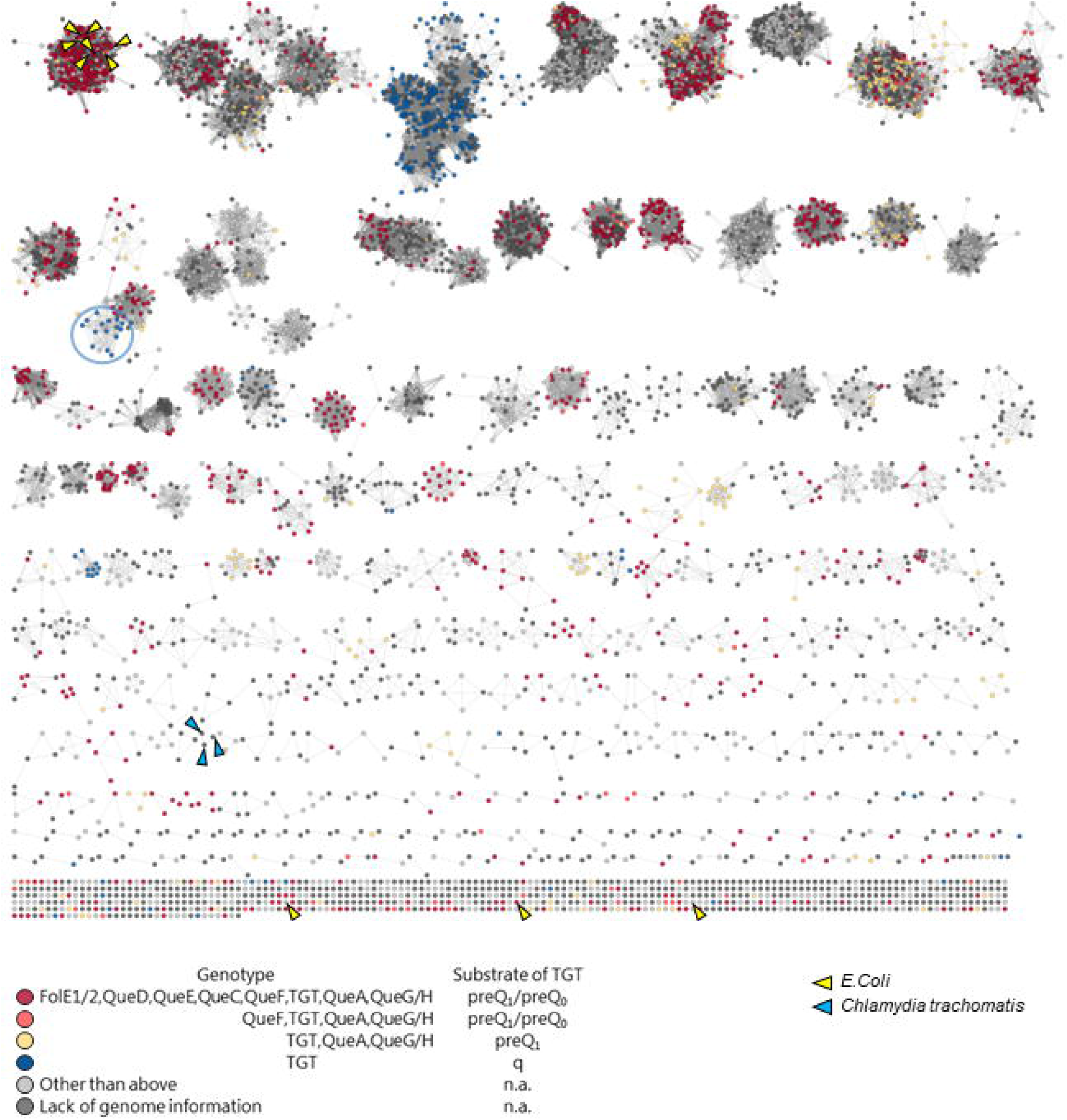
Protein sequence similarity network (SSN) of 7,625 QPTR (PF02592 family) proteins. Each node in the network represents a QPTR protein. An edge (represented as a line) is drawn between two nodes with a BLAST *E*-value cutoff of better than 10^−70^ (alignment score of 70). The nodes are colored based on the presence/absence of the other Q synthesis genes in the corresponding genome. Cases with inconclusive Q pathway gene distribution are colored in light gray. QPTR proteins without genome information are colored in dark gray. QPTR homologs from *E. coli* and *C. trachomatis* are indicated by yellow and blue arrows, respectively. The solitary TGTs from genomes harboring only *tgt* that cluster with TGTs in complete Q pathways are circled in blue.

Following up on this discrepancy, we retrieved all Q synthesis proteins from the InterPro database with their IPR protein family ID (see Materials and Methods) and counted the numbers of each protein encoded in each genome at the level of different taxonomic ranks, including order, class, and family (Table S1). Then, we extracted the TGT sequences from all orders, classes, families, and genera that encode only *tgt* as described in the methods section, aligned them, and compared the predicted substrate-binding residues (Table S2). As we previously reported (15), TGTs that salvage q typically contains GG[LS][AS]G in the substrate-binding pocket (Fig. 3). Interestingly, bTGTs in *Bartonella* and *Pelagibacter* contain a GGLAVG site like the *E. coli* enzyme (Fig. 3) and are phylogenetically distant from other bTGTs found in genomes that only encode *tgt* (Fig. 3). We then examined the substrate-binding pocket in a modeled structure of *B*. *henselae* bTGT (Bh bTGT, UniProt ID A0A0R4J8M4_BARHE) aligned with the structure of the human TGT catalytic subunit QTRT1 in complex with q (PDB ID: 6H45) (22). The aspartate residues and G216GLAVGE222 that are conserved in bTGT proteins of the *Hyphomicrobiales* order are in proximity to queuine (Fig. S1B and C). The predicted distance between V220 and the cis-diol groups of q is less than 1 Angstrom (Fig. S1C), suggesting that it may prevent the binding of q to *B. henselae* bTGT as in the *Zymomonas mobilis* TGT (23). We previously showed that *C. trachomatis* TGT salvages q and that its substrate-binding pocket can accommodate the larger substrate (15). Here, we propose that the bTGT protein and the QPTR transporter in *Bartonella* and *Pelagibacter* species salvage preQ_1_, even though they lack, like *C. trachomatis*, the enzymes that make Q-tRNAs from preQ_1_-tRNAs.

**Figure 3.**
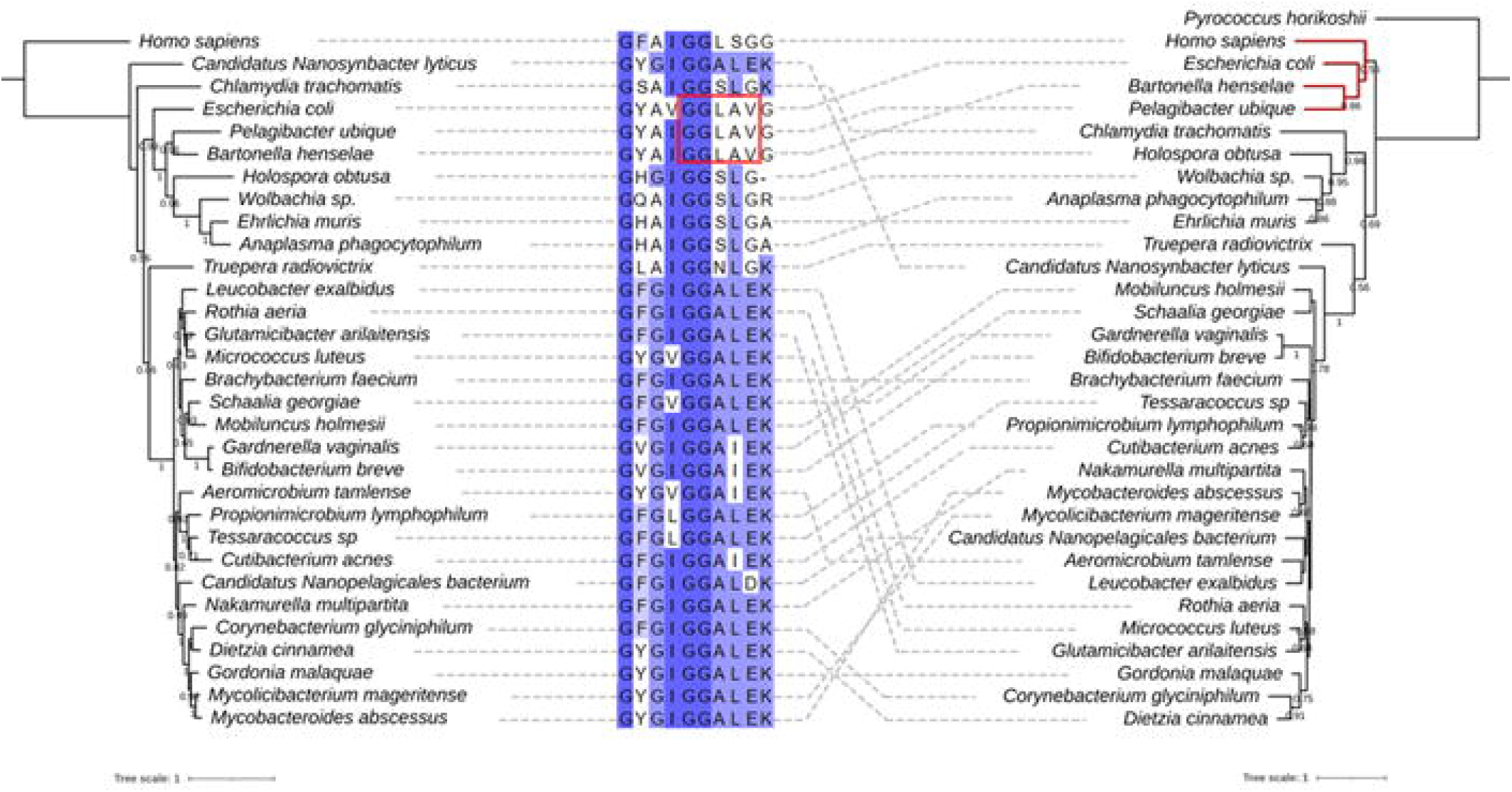
Comparison of the phylogenetic tree of taxonomic ranks of bacteria that harbor *tgt* only and the maximum likelihood tree of their TGT proteins. The maximum likelihood tree of marker proteins of taxonomic ranks of bacteria that harbor *tgt* only without other Q proteins (left). The maximum likelihood tree of TGT proteins from organisms of each corresponding rank (right). Human and *Pyrococcus horikoshii* tRNA-guanine (15) transglycosylases were used as the outgroup for each tree, respectively. TGT of *E. coli* was used for comparison. Bootstraps above 0.5 are shown under the branch. A multiple sequence alignment of consensus residues in the TGT proteins in the moiety of 7-substitue of deazapurine is shown (middle). The residues are highlighted by the percent identity. The dashed lines connect each organism and their TGT sequence. The branch of TGTs from *Bartonella, Pelagibacter, E. coli*, and humans is highlighted in red.

### *B. henselae* bTGT and QPTR proteins preferentially salvage preQ_1_ but also q and Q with low affinity

The gene encoding the *B. henselae* Bh bTGT protein complemented the Q-phenotype of a *queDF tgt* deletion mutant of *E. coli* when expressed *in trans* and in the presence of exogenous preQ_1_ even in low concentrations (down to 10 nM) (Fig. 4A). Similarly, Bh QPTR transported both preQ_1_ and preQ_0_ in an *E. coli* strain auxotrophic for preQ_1_ and preQ_0_ (Fig. 4B). These results confirmed our predictions based on the SSN that the *Bartonella* bTGT and QPTR proteins use preQ_1_ as a substrate but do not match the metabolic reconstruction that predicted q salvage in *B. henselae*. Hence, we tested if this organism’s bTGT and QPTR could have evolved a broader substrate specificity and use q as a substrate.

**Figure 4.**
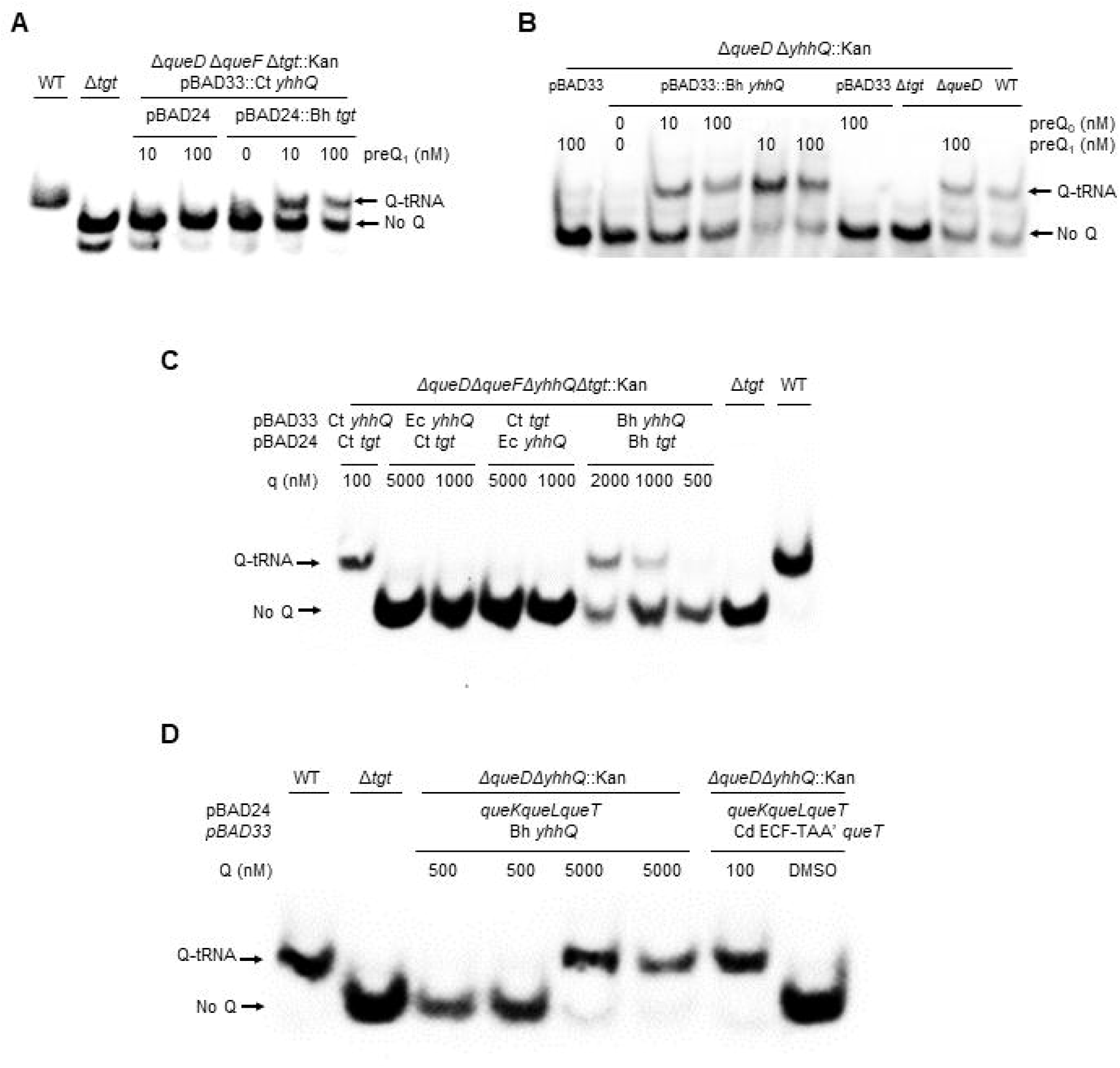
Bh TGT and Bh YhhQ salvage preQ_1_ in *E. coli*. Detection of Q-tRNAAsp GUC by the APB assay. Q-modified tRNAs that migrate slower are indicated by an arrow. tRNAs were extracted from WT and mutant strains expressing different Q salvage genes. The strains used are denoted in the first line. The genes and corresponding vectors are indicated in the second line. Plasmid and strain information are given in Table S9. Cells were grown in minimal media in the presence of exogenous preQ_0_, preQ_1_, q or Q as noted. DMSO was used as control when no deazapurine was supplemented.

To test whether the QPTR and bTGT proteins of *B. henselae* can use q as a substrate, we used an *E. coli* strain that expresses the QPTR and/or the bTGT proteins of *C. trachomatis* that can only use q as a substrate (Fig. 1) (15). Salvage of q was observed when expressing both Bh *tgt* and *yhhQ* genes at concentrations of q over 500 nM (Fig. 4C). When overexpressing the *E. coli yhhQ* gene, no such salvage of q was observed even at concentrations of 5 μM (Fig. 4C). However, if the *C. trachomatis* bTGT and QPTR proteins can salvage q when present at 100 nM, the Bh QPTR cannot (Fig. S2). These results showed that the Bh QPTR protein has acquired the capacity to transport q while retaining the preQ_1_/preQ_0_ specificity, but it is still not as efficient as the *C. trachomatis* QPTR transporter that is specific for q.

We then tested whether the Bh QPTR could transport the Q ribonucleoside by modeling with an *E. coli* strain expressing the *C. difficile* Q hydrolase (Cd *queK*) and q lyase (Cd *queL*) genes that allow Q to be salvaged by *E. coli* (Fig. 1) (15). Expressing only Bh QPTR allowed the salvage of Q only at extremely high concentrations (5 μM) (Fig. 4D). When the *C. difficile* Q transporter (Cd ECF_TAA’, QueT) is expressed in this strain, Q can be salvaged when present at concentrations of 100 nM [(15) and (Fig. 4D].

### Traces of preQ_1_ can be detected in endogenous *B. henselae* tRNAs

The natural habitat of this intracellular pathogen (mammalian vasculature) should be richer in q than in preQ_1_. We, therefore, analyzed by LC-MS/MS the tRNA modification profile of bulk tRNA extracted from *B. henselae* cells grown in sheep blood agar (HIBB) medium. The experiments were performed three times independently with conflicting results. The first two experiments were done to test the tRNA extraction protocols with an intracellular bacterium, using only one sample each time. Small amounts of preQ_1_ and minute amounts of Q were detected the first time (but neither was present the second time (data not shown). Because cells were grown in the presence of sheep blood and serum, we had little control over the sources of Q or preQ_1_; we, therefore, repeated the experiment a third time with 5 independent samples, adding 100 nM preQ_1_ in three of the samples. As shown in Fig. 5, Fig. S3 and Table S3, small amounts of preQ_1_ and Q were detected in all samples (around 1000 times less than in a typical *E. coli* sample). The exogenous addition of 100 nM preQ_1_ did not make any difference. It is not possible to determine if the observed Q was derived from *B. henselae* tRNAs or from contaminating mammalian tRNAs, as eukaryotic-specific tRNA modifications such as m^2^_2_G, are detected in similar quantities (Table S3) and great variations in Q levels were observed between samples. Nevertheless, the presence of preQ_1_ cannot be explained by any contamination as mammalian host tRNAs never harbor this modification and other bacteria would not accumulate it, as preQ_1_ had only been detected previously in *queA* mutants of *E. coli* (24). The low amounts detected suggest that the tRNAs are not fully modified, as the preQ_1_ levels are 11%-40% of cmnm^5^s^2^U levels and 1.1%-2.8% of k^2^C levels; two well-conserved bacterial modifications (25)(Fig. S4 and Table S3).

**Figure 5.**
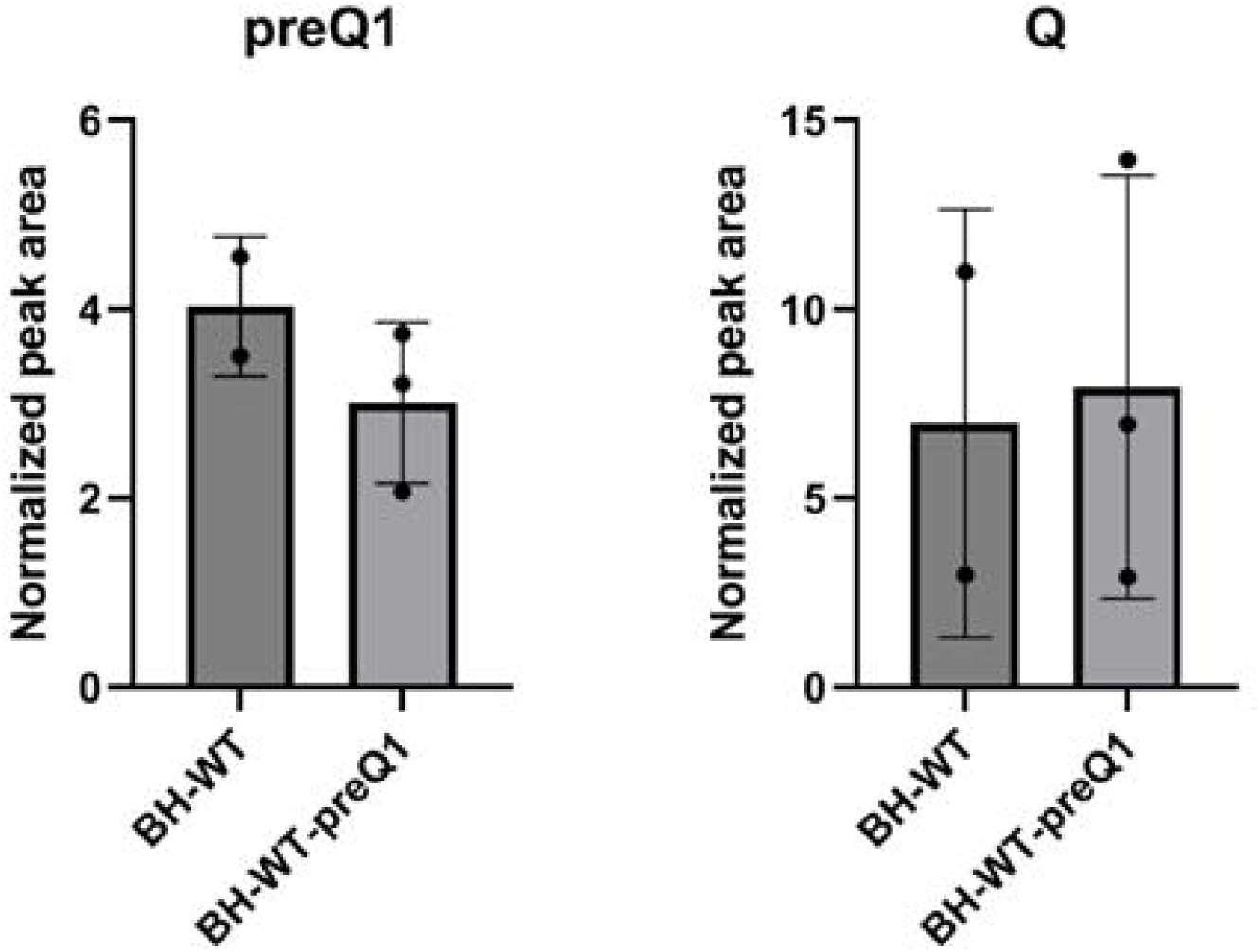
Quantification of preQ1 and Q in *B. henselae*. LC-MS/MS analysis of ribonucleosides containing preQ1 and Q was performed on 600 ng of hydrolyzed small RNA, with signal intensities normalized to the sum of UV absorbances of the canonical ribonucleosides, using an inline UV detector, to correct for differences in amounts of injected RNA. Data represent mean ± SD for three biological replicates of *B. henselae* BH-WT+preQ1 (BH-WT+preQ1) and mean ± deviation about the mean for two biological replicates of *B. henselae* BH-WT (BH-WT); technical duplicates were performed for each biological replicate. All modifications were validated with standards.

### Early loss of Q pathway genes in the *Bartonellaceae* family within the *Hyphomicrobiales* order

*B. henselae* belongs to the Alphaproteobacteria class (25). This is a diverse Gram-negative taxon comprised of several phototrophic genera, several genera metabolizing C1-compounds (e.g., *Methylobacterium* spp.), symbionts of plants (e.g., *Rhizobium* spp.), endosymbionts of arthropods (*Wolbachia*) and intracellular pathogens (e.g., *Rickettsia*) (26). To better understand the evolution of the Q synthesis and salvage pathway in Alphaproteobacteria, we performed a phylogenetic distribution analysis of the corresponding genes in 2,127 different species with complete genome sequences from the class in the BV-BRC database as described in the methods section (Fig. S5). The tree suggests that there were three events involving the loss of preQ_1_ synthesis genes: one occurred after the split between *Brucella* and *Bartonella* species (Fig. 6 and Fig. S5 node 1), one occurred after the split of *Pelagibacteraceae* (Fig. S5 and Fig S6 node 2), and one occurred after the split of *Anaplasmataceae* (Fig. S5 and Fig S6 node 3). This analysis revealed that most Alphaproteobacteria are prototrophic for Q, as 1,414 (64%) of the species analyzed encode the complete pathway. In addition, if the loss of the preQ_1_ synthesis genes occurred sporadically in different branches, all the organisms analyzed, with the exception of *Bartonella,* harbored *tgt*, *queA*, and *queG* genes and hence were not predicted to salvage q. Focusing more specifically on the *Bartonella* genus using a similar analysis revealed a very different pattern (Fig. 6).

**Figure 6.**
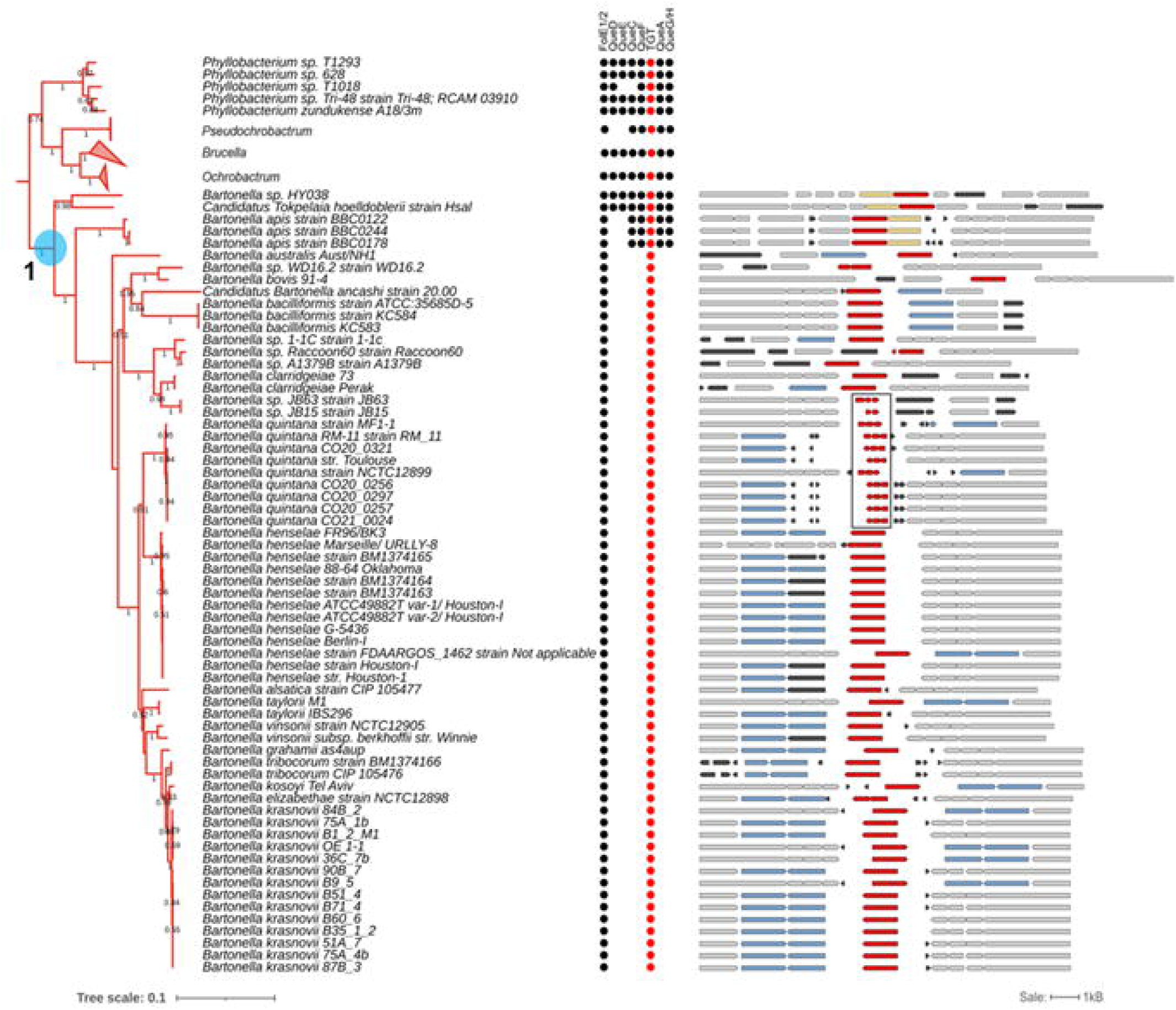
Phylogenetic analysis of Queuosine biosynthesis genes in *Bartonellaceae*. The clade of *Bartonella spp.* (*Bartonellaceae*; Alphaproteobacteria; left). The branches of *Pseudochrobactrum, Brucella*, and *Ochrobactrum* were collapsed. The highlighted node 1 corresponds to the node in the tree in Figure S5. The presence of Q biosynthesis proteins is indicated by the circles with TGTs highlighted in red. The comparative view of the corresponding truncated and full-length *tgt* gene variants in different *Bartonella* species in the tree (right). Red, *tgt*; yellow, *queA*; blue, genes encoding a transporter-like protein; black, hypothetical genes; gray, other genes. The fragmented *tgt* genes in *B. quintana* strains are boxed. Gene IDs are provided in supplementary Table S4.

Among the 65 organisms in the *Bartonella* genus with available complete genomes, as of October 2023, only *Bartonella* sp. HY038, branching at the root of the genus, encoded the canonical *de novo* Q synthesis pathway. Most, like *B. henselae*, have lost all the genes but *tgt*. In addition, a more in-depth analysis revealed fragmented *tgt* genes of several *B. quintana* strains (boxed in Fig. 6), suggesting these organisms have totally lost the capacity to make Q-modified tRNAs. This evolutionary scenario seems to be a recurring theme in intracellular bacteria like the *Rickettsiales*. While nearly all rickettsiae branching closer to the root retain the full pathway except for *queD* (collapsed in Fig. S6), other rickettsiae such as *Anaplasma, Ehrlichia,* and *Wolbachia*, have lost nearly the full pathway. However, cases of fragmented *tgt* genes are rare and only observed in a *Wolbachia* endosymbiont of *Cimex lectularius* (box in Fig. S5).

In summary, the phylogenic distribution analysis suggests that the direct ancestor of *Bartonella* species must have harbored the full Q pathway but that it was lost very early in the evolution of this clade. Most bacteria in this clade are predicted to transport and insert preQ_1_ but without further conversion to Q. In addition, some species like *B. quintana* have lost the pathway completely (Fig. 6).

## Discussion

Q is an ancient modification predicted to be present in the ancestors of bacteria (27). It is still present in most extant bacteria even if minimalist genetic codes can exist without this complex modification (28). Independent analyses of the genomes of bacteria in the human microbiome have shown that 90 to 95% of these organisms maintain the capacity to synthesize or salvage Q with around half encoding the full synthesis pathway (15,29). Many of the bacteria that have lost Q are organisms that have undergone a genome reduction process, where their genetic material has been streamlined over evolutionary time such as the parasitic *Mycoplasma* spp. or insect endosymbionts such as *Riesia pediculicola* (30,31).

Obligate intracellular human pathogens tend to have reduced genomes compared to their free-living ancestors as their metabolisms have adapted to a nutritionally rich niche (32). Regarding Q, the scenarios that one can envision in the transition to a strict intracellular lifestyle with access to the queuine precursor from the mammalian host are: 1) keeping the ancestral pathway; 2) losing the modification; or 3) switching to a queuine salvage route. We performed the metabolic reconstruction of Q metabolism in genera of strict intracellular human pathogens such as *Chlamydia* spp., members of the order Rickettsiales (*Anaplasma* spp., and *Rickettsia* spp.) and *Coxiella burnetii* (33,34). Indeed, examples of these three possible paths were observed (Fig. 7): *Rickettsia* (Fig. 7A) and *Coxiella* (Table S5 line 267) have kept the full Q synthesis pathway; *Anaplasma* spp. (Fig. 7A), *Borrelia* (Fig. 7B), *Ehrlichia* spp. (Table S5 line 52), and *Chlamydia* (Table S5 line 212), kept only *tgt*. Most species in Mycobacteriales have totally lost the pathway genes except *tgt* (Fig. 7C).

**Figure 7.**
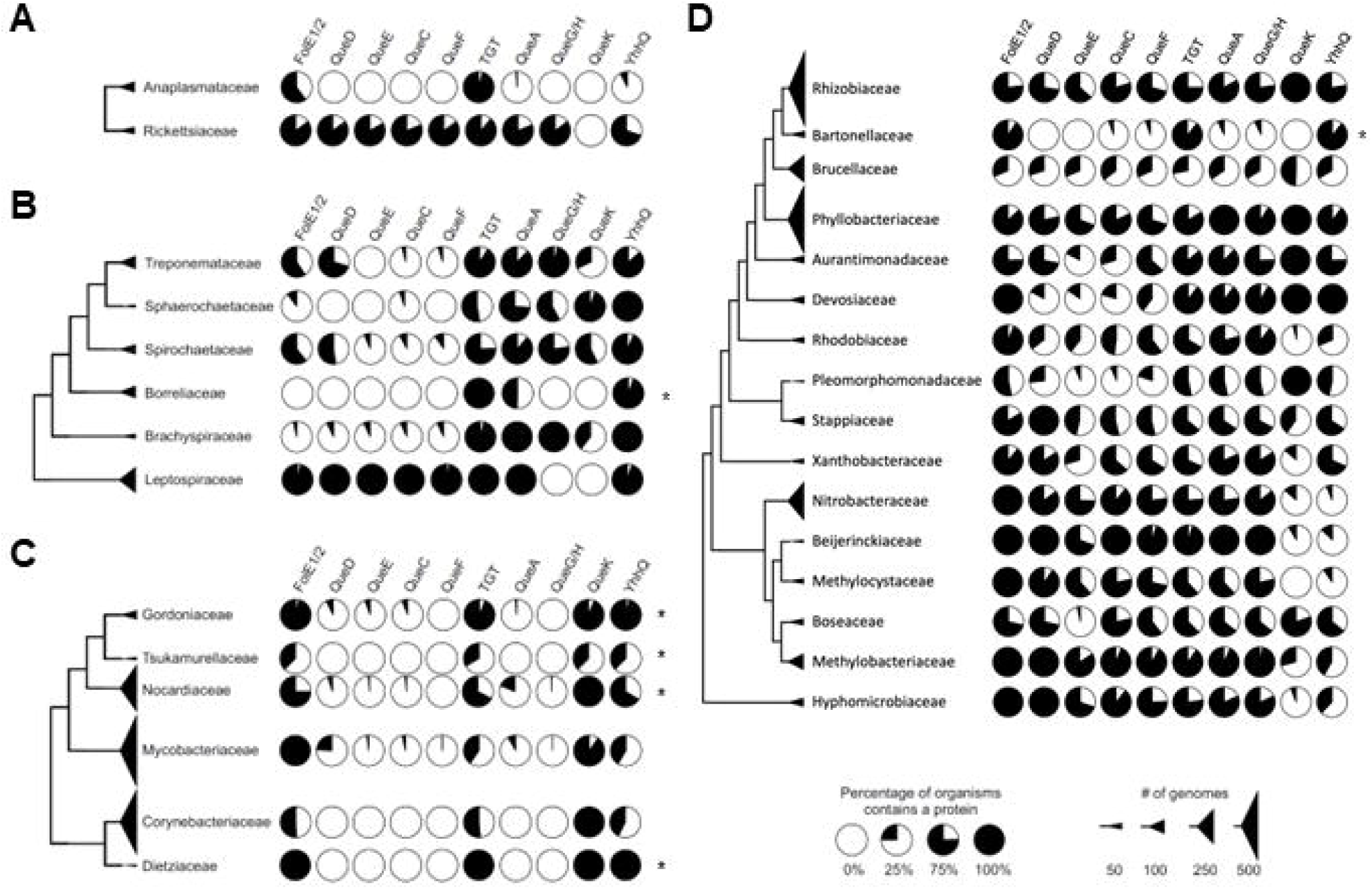
The prevalence of Q pathway proteins in selected taxonomic ranks in Uniprot database. (A) A cladogram of the order Hyphomicrobiales. (B) A cladogram of the order Mycobacteriales. (C) A cladogram of the class Spirochaetia. (D) A cladogram of the order Rickettsiales. Each pie chart represents the percentage of organisms in each taxonomic rank that contains the corresponding Q pathway protein. The size of each triangle correlates with the number of genomes in each taxonomic unit. The sequences of TGTs’ substrate binding sites from ranks that encoded only *tgt* were analyzed and presented in Table. S2. Only taxonomic ranks that contain more than 15 TGTs are shown.

The situation seen in *B. henselae* is not commonly observed in other intracellular bacteria and no other organisms in the Hyphomicrobiales order seem to follow the same pattern (Fig. 7D). Indeed, the presence of bTGT and QPTR homologs and the absence of QueA and QueG or QueH (Fig. 6) would suggest that these organisms salvage q like *C. trachomatis*, but the corresponding enzymes have retained their substrate specificity towards preQ_1_ (Figs. 3 and 4). PreQ_1_ is not a molecule found in mammalian cells (Brian Green, personal communication), and the fact that we were able to detect a small proportion of tRNAs carrying that modification when extracted from *B. henselae* cells grown in HIBB suggests this pathway is functional even if the source of preQ_1_ remains a mystery and could be due to contamination with a preQ_1_ synthesizing organism, an unknown source of preQ_1_ in the culture medium, or from the activity of a yet-to-be-discovered q hydrolase (intracellular or possibly extracellular as q transport is not efficient as discussed below) (Fig. 1D). The low amount of preQ_1_ modification (Table S3) suggests it does not play an important role in decoding accuracy under these conditions. The fact that *tgt* gene decay is observed in several organisms in this clade such as *B. quintana* reinforces this idea. A primary difference between *B. quintana* and *B. henselae* is their reservoir ecology. *B. quintana* uses only humans as a reservoir, whereas *B. henselae* is more promiscuous and frequently isolated from both cats and humans (35).

Another intriguing finding of this study is that the QPTR and bTGT enzymes of *B. henselae* can use q as a substrate when present at high concentrations (>500 nM), whereas the *E. coli* orthologs cannot. It is difficult to establish if such concentrations could be physiological, and we could not show with certainty that *B. henselae* tRNA extracted from HIBB-grown cells harbored Q. One can propose several evolutionary explanations for this broadening of substrate specificity of the *B. henselae* salvage proteins. In one, the enzyme and transporter specificities would become more relaxed in *B. henselae* as they are no longer under selective pressure to maintain efficient preQ_1_ selection, and we are observing an intermediate stage along the evolutionary loss of the whole pathway. An argument against that hypothesis is that if preQ_1_ synthesis genes and *queA*, *queG* and *queH* genes seem to have been lost very early in the clade, *tgt* is often the last maintained and we did not find any examples where *tgt* was lost with the other genes maintained, suggesting a fitness advantage of maintaining the *tgt* gene. In addition, both *B. henselae* QPTR and bTGT encoding genes are expressed and the proteins detected *in vitro* [see Table S5 of (36)]. The other possibility is that we are observing a transition of a preQ_1_ salvage to a q salvage that is working poorly in human cells but could be efficient in an environment with more q/Q. Could the insect vector provide such an environment? Answering these questions will require further studies, including additional quantitative data on q/preQ_1_ levels in different environments and tRNA modification profiles along the pathogen’s life cycle. In summary, the study sheds light on the diverse and adaptable nature of queuosine metabolism in various bacteria, particularly in intracellular pathogens. The unique characteristics of Q salvage observed in *B. henselae* raise intriguing questions about its role in different host/vector environments. Further investigations are warranted to unravel the complexities of Q salvage and its implications to *Bartonella*’s virulence.

## Materials and methods

### Comparative genomics and bioinformatics

The BLAST tools (37) resources at the National Center for Biotechnology Information (NCBI) and BV-BRC (38) were routinely used. Multiple sequence alignments were built using MUSCLE v5. 1 (39) and visualized and edited with Jalview2 (40). Protein sequences were retrieved from the NCBI using the following accession numbers: Ct YhhQ, NP_219643.1; Ct TGT, NP_219697.1; Cd ECF-A, YP_001086568.1; Cd ECF-A, YP_001086569.1; Cd ECF-T, YP_001086570.1; Cd Queuosine hydrolase, YP_001088185.1; Cd ECF-S, YP_001088186.1; Cd Queuine lyase, YP_001088187.1; Ec YhhQ, WP 001100469.1; Ec bTGT; Bh YhhQ, WP_011181356.; Bh bTGT, WP_011180873.1. Protein IDs used for this study are listed in Table S6. Complete genomes of Alphaproteobacteria with good quality were retrieved from BV-BRC (38).

For metabolic reconstruction analyses in each taxonomic rank, all protein members were retrieved from the InterPro database (41) using the following IPR family ID: FolE1, IPR001474; FolE2, IPR003801; QueD, IPR007115; QueE, IPR024924; QueC, IPR018317; QueF, IPR00029500; TGT, IPR004803; QueA, IPR003699; QueG, IPR004453; QueH, IPR003828. A universal single-copy small ribosomal protein uS2, (IPR001865) was used to estimate the number of organisms in each rank. The number of proteins per taxonomic rank was computed and the criteria used for filtering the groups ranks encoding just TGT were the following: 1) the number of TGT proteins was no less than 10 so we were not polluted with small taxonomic sample size; 2) the number of each of the QueDECFAGH proteins was no more than a fifth of the number of TGT proteins. To analyze the conserved residues in the substrate-binding pockets, the sequences of TGT proteins from select taxonomic groups were retrieved from UniProt and aligned using MUSCLE v5.1 (39). The conserved residues were visualized using weblogo3 (42).

The structure of TGT proteins (*Bartonella henselae* str. Houston-1 bTGT, A0A0R4J8M4; *Anaplasma phagocytophilum* bTGT, S6G6J1; *Nakamurella multipartita* bTGT, C8X7A7) were modeled using SWISS-MODEL (40) using Alpha Fold structure (*Bartonella fuyuanensis* bTGT (ID A0A840DZ06_9HYPH), *Anaplasma phagocytophilum* str ApMUC09 TGT (ID A0A0F3NAG3_ANAPH), and *Nakamurella multipartita* bTGT (ID C8X7A7_NAKMY) as templates respectively (43). The cartoon representation of protein structure was produced by PyMol (version 2.5) (44) and colored by domain (red, N-terminus; light blue, C-terminus).

### Sequence Similarity Networks (SSNs)

The Enzyme Function Initiative (EFI) suite of web tools was used to generate the SSN (45). Visualization of SSNs was carried out using Cytoscape 3.10.1ape (46). 7,625 PF02592 family sequences were retrieved from UniProt using the family option with fraction of 3 and submitted to EFI. The initial SSN was generated with an alignment score cutoff set such that each connection (edge) represents a sequence identity of above approximately 40%. The obtained SSN was first colored according to the configurations for salvaging preQ_1_, preQ_0_, queuine, and Queuosine *de novo* synthesis. Then more stringent SSNs were created by increasing the alignment score cutoff in small increments (usually by 5). This process was repeated until most clusters were homogeneous in their colors. The UniProt IDs were associated with the genome ID including GenBank/EMBL, RefSeq nucleotide, BV-BRC genome ID, Ensembl genome ID, using homemade scripts (scripts available upon request). The UniProt IDs of PF02592 family sequences in the SSN are listed in Table S7 as well as corresponding presence of Q pathway genes. The connection between UniProt IDs and genome information was performed by querying UniProt ID mapping file using homemade scripts (scripts available upon request).

### Phylogenetic investigations and Q gene presence/absence distribution pattern analysis

For phylogenetic analysis of species, the 20 ribosomal protein data (L2, L3, L4, L5, L6, L9, L10, L14, L16, L18, L22, L24, S2, S3, S4, S5, S8, S10, S17, and S19) were retrieved from BV-BRC and aligned independently using MUSCLE v5.1 (39). Alignments were trimmed using BMGE v1.12 using default settings (47) and then concatenated. To build the phylogenetic tree of TGT proteins, the sequences of TGT proteins from each taxonomic rank were retrieved (ID are listed in Table S6) and subject to alignment with MUSCLE (39) and trimming with BMGE (47). The maximum likelihood tree was built by FastTree (48) using the LG-cat model with a hundred bootstraps. Trees were visualized using iTOL (49). To view the gene clustering near *tgt* in different bacterial genomes, GizmoGene (http://www.gizmogene.com) was used to extract gene regions and subjected to visualization using Gene Graphics (50). The genomic regions are listed in Table S4.

### Strains, media, and growth conditions

Strains and plasmids used in this study are listed in Table S9. LB medium (tryptone 10 grams/liter, yeast extract 5 grams/liter, sodium chloride 5 grams/liter) was routinely used for *E. coli* growth at 37°C. The medium was solidified using 15 g/L of agar. As needed, kanamycin (50 g/mL), ampicillin (100 g/mL), and chloramphenicol (25 g/mL) were added. In the presence of exogenous Q precursors as previously described (51), cells were cultured in M9-defined medium containing 1% glycerol (Thermo Fisher Scientific, Waltham, MA, USA) for the purpose of eliminating background Q-tRNA. After cells reached an optical density at 600 nm (OD_600nm_) of 0.1-0.2, 0.2% arabinose was added to induce the expression of genes under the pBAD promoter. After cells reached an OD_600nm_ of 0.2, DMSO, preQ_0_, preQ_1_, q, or Q were added. The transport reaction was stopped at time points of 30 or 60 min after supplementing with DMSO or different Q precursors by placing samples on melting ice and then centrifuging, followed by tRNA extraction. Q was purchased from Epitoire (Singapore), q from Santa Cruz Biotechnology, preQ_1_ and preQ_0_ from Sigma-Aldrich.

*B. henselae* Houston I was obtained from the American Type Culture Collection (ATCC 49882) and cultivated as previously described (52) on HIBB agar plates [Bacto heart infusion agar (Becton, Dickinson, Sparks, MD) supplemented with 4% defibrinated sheep blood and 2% sheep serum (Quad Five, Ryegate, MT) by volume] for 4 days at 37°C, 5% CO_2_ and 100% relative humidity. When required, preQ_1_ was added to a final concentration of 100 nM. Following harvest into ice-cold heart infusion broth, tRNA was collected from the bacterial cells.

### Construction of *E. coli* strains and plasmids

*B. henselae yhhQ* gene (Bh *yhhQ*) was chemically synthesized (without optimization) in pTWIST^_^Kan vector. Xbal and HindIII restrictions sites were added at the 5′ and 3′ ends, respectively (Twist Bioscience HQ) (**Table S10**). Bh *yhhQ* DNA sequence was amplified using two primers pairs (F_Bh yhhQ_XbaI_PBAD33 and R_Bh yhhQ_HindIII_PBAD33) by PCR with the addition of restrictions sites XbaI and HindIII at their 5′ and 3′ ends, respectively. Bh *yhhQ* was cloned into the XbaI and HindIII sites of pBAD33. *B. henselae tgt* (Bh *tgt*) was amplified by PCR from *B. henselae* genomic DNA using the KpnI-RBS-TGTBh-F_PBAD24 and TGTBh-SbfI-R _PBAD24 primers and cloned into the KpnI and SbfI sites of pBAD24. The UniProt IDs for Bh *yhhQ* and Bh *tgt* are A0A0H3M726_BARHE and A0A0R4J8M4_BARHE, respectively. *E. coli* transformations were performed using the CaCl_2_ chemical transformation procedure (53). Transformants were selected on LB agar supplemented with ampicillin or chloramphenicol (100 µg/mL). The clones were validated through sequencing and PCR analyses using primers designed specifically for Bh *yhhQ* and Bh *tgt* genes. All primers used in this study are listed in Table S10.

### tRNA extraction and migration

Cells were harvested by centrifugation at 16,000 x g for 2 minutes at 4°C. Immediately after pelleting, the cells were resuspended in 1 mL of Trizol (Thermo Fisher Scientific, Waltham, MA, USA). According to the manufacturer’s instructions, small RNA was extracted with the PureLink^Tm^ miRNA Isolation kit (Thermo Fisher Scientific, Waltham, MA, USA). 50 μL of RNase-free water were used to elute the purified RNAs. Quantification of prepared tRNA was performed using a Nanodrop 1000 spectrophotometer. We loaded 150 ng of tRNAs per well on a denaturing 8 M urea, 8% polyacrylamide gel containing 0.5% 3- (Acrylamido) phenylboronic acid (Sigma-Aldrich) after resuspending in a 2X RNA Loading Dye (NEB). Migration was performed in a mixture of 1X TAE at 4°C. With a wet transfer apparatus in 1X TAE at 150 mA at 4°C for 90 minutes, tRNAs were transferred onto a Biodyne B precut nylon membrane (Thermo Scientific). The membrane was UV irradiated in a UV crosslinker (Fisher FB-UVXL-1000) at a preset UV energy dosage of 120 mJ/cm2. The North2South Chemiluminescent Hybridization and Detection Kit (Thermo) was used to detect tRNA^Asp^. As the DIG Easy Hyb (Roche) drastically reduces the background noise, it was used as the initial membrane-blocking buffer instead of the North2South kit’s membrane-blocking buffer. Hybridization was done at 60°C, using the specific biotinylated primer for tRNA Asp GUC (14) (5’ biotin-CCCTGCGTGACAGGCAGG 3’ for *E. coli* added to a final concentration of 50 ng/mL. The blot was visualized by the iBright™ Imaging Systems.

### tRNA profiling by mass spectrometry

tRNA for each sample (1.8 µg) was hydrolyzed in a 30 µL digestion cocktail containing 2.49 U benzonase, 3 U CIAP (calf intestinal alkaline phosphatase), 0.07 U PDE I (phosphodiesterase I), 0.1 mM deferoxamine, 0.1 mM BHT (butylated hydroxytoluene), 3 ng coformycin, 25 nM 15N-dA (internal standard [^15^N]_5_- deoxyadenosine), 2.5 mM MgCl_2_ and 5 mM Tris-HCL buffer pH 8.0. The digestion mixture was incubated at 37 °C for 6 h. After digestion, all samples were analyzed by chromatography-coupled triple-quadrupole mass spectrometry (LC-MS/MS). For each sample, 600 ng of hydrolysate was injected for two technical replicates. Using synthetic standards, HPLC retention times of RNA modifications were confirmed on a Waters Acuity BEH C18 column (50 × 2.1 mm inner diameter, 1.7 µm particle size) coupled to an Agilent 1290 HPLC system and an Agilent 6495 triple-quad mass spectrometer. The Agilent sample vial insert was used. The HPLC system was operated at 25 °C and a flow rate of 0.3 mL/min in a gradient Table S11 with Buffer A (0.02% formic acid in double distilled water) and Buffer B (0.02% formic acid in 70% acetonitrile). The HPLC column was coupled to the mass spectrometer with an electrospray ionization source in positive mode with the following parameters: Dry gas temperature, 200 °C; gas flow, 11 L/min; nebulizer, 20 psi; sheath gas temperature, 300 °C; sheath gas flow, 12 L/min; capillary voltage, 3000 V; nozzle voltage, 0 V. Multiple reaction monitoring (MRM) mode was used for detection of product ions derived from the precursor ions for all the RNA modifications with instrument parameters including the collision energy (CE) optimized for maximal sensitivity for the modification. Based on synthetic standards (Biosynth) with optimized collision energies, the following transitions and retention times (except k^2^C, which we do not have standard) were monitored: cmnm^5^s^2^U, m/z 348 → 141, 2.36 min; k^2^C, m/z 372.1 → 240.1; m^2,2^G, m/z 312 → 180, 9.70 min; preQ_1_, m/z 312 → 163, 2.15 min; Q, m/z 410 → 163, 5.53 min. Signal intensities for each ribonucleoside were normalized by dividing by the sum of the UV signal intensities of the four canonical ribonucleosides recorded with an in-line UV spectrophotometer at 260 nm.

## Supporting information

SupTablesS1toS11

FiguresS1toS6

## Acknowledgments

We are thankful to Dr Brian Greene for communicating unpublished results.

