## Supplementary material for "Queuosine Salvage in *Bartonella henselae* Houston 1: A Unique Evolutionary Path": FiguresS1toS6

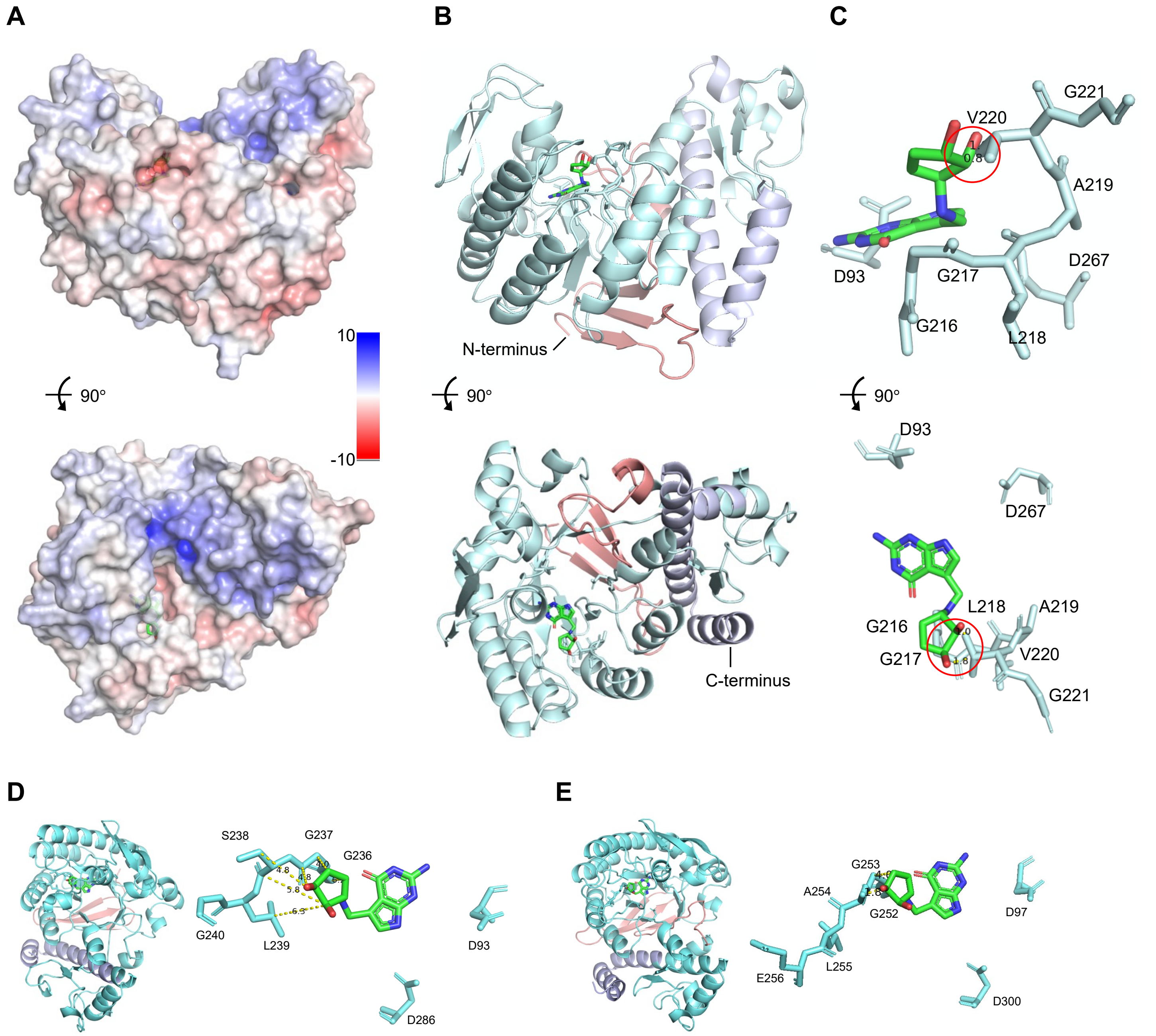


**Figure S1. The predicted structure of Bh TGT.** (A) Electrostatic potential mapped on the surface of Bh TGT, where positive charges are shown in blue, negative charges in red, and neutral charges in white. Queuine is shown in stick format and colored by atom type. (B) Structure of Bh TGT. The cartoon representation was colored by domain. The stick cartoon represents the key residues in the proximity of queuine (C). The distance between queuine and V220 is 0.8~1.8 Angstrom (red circle). Predicted structures of *Anaplasma phagocytophilum* TGT (Uniprot S6G6J1) (D) and *Nakamurella multipartita* TGT (C8X7A7) (E). The cartoon representation was colored by domain. The stick cartoons represent the key residues in the proximity of queuine. The distance between queuine and GGSLG and GGALE above is 4.0 and 1.8 Angstrom, respectively.

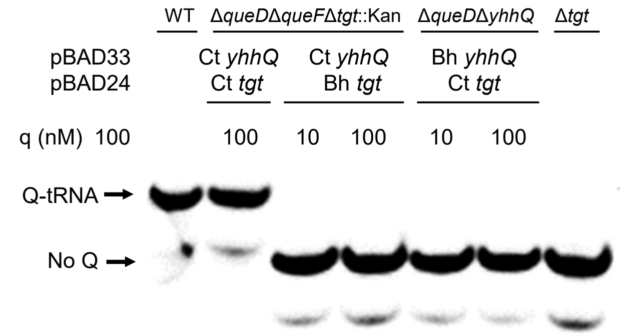


**Figure S2. Bh YhhQ and Bh TGT do not salvage q as efficiently as Ct YhhQ and Ct TGT .** Detection of Q-tRNA$^{Asp}{}_{GUC}$ by the APB assay, Q-modified tRNAs that migrate slower are indicated by an arrow. tRNAs were extracted from WT or different Q$^{-}$ *E. coli* mutant expressing the *yhhQ* and *tgt* genes from Bh and Ct, as indicated in the figure, in a minimal media in the presence of exogenous q. The strains used are denoted in the first line. The genes and corresponding vectors are indicated in the second line. Plasmid and strain information are given in Table S9.

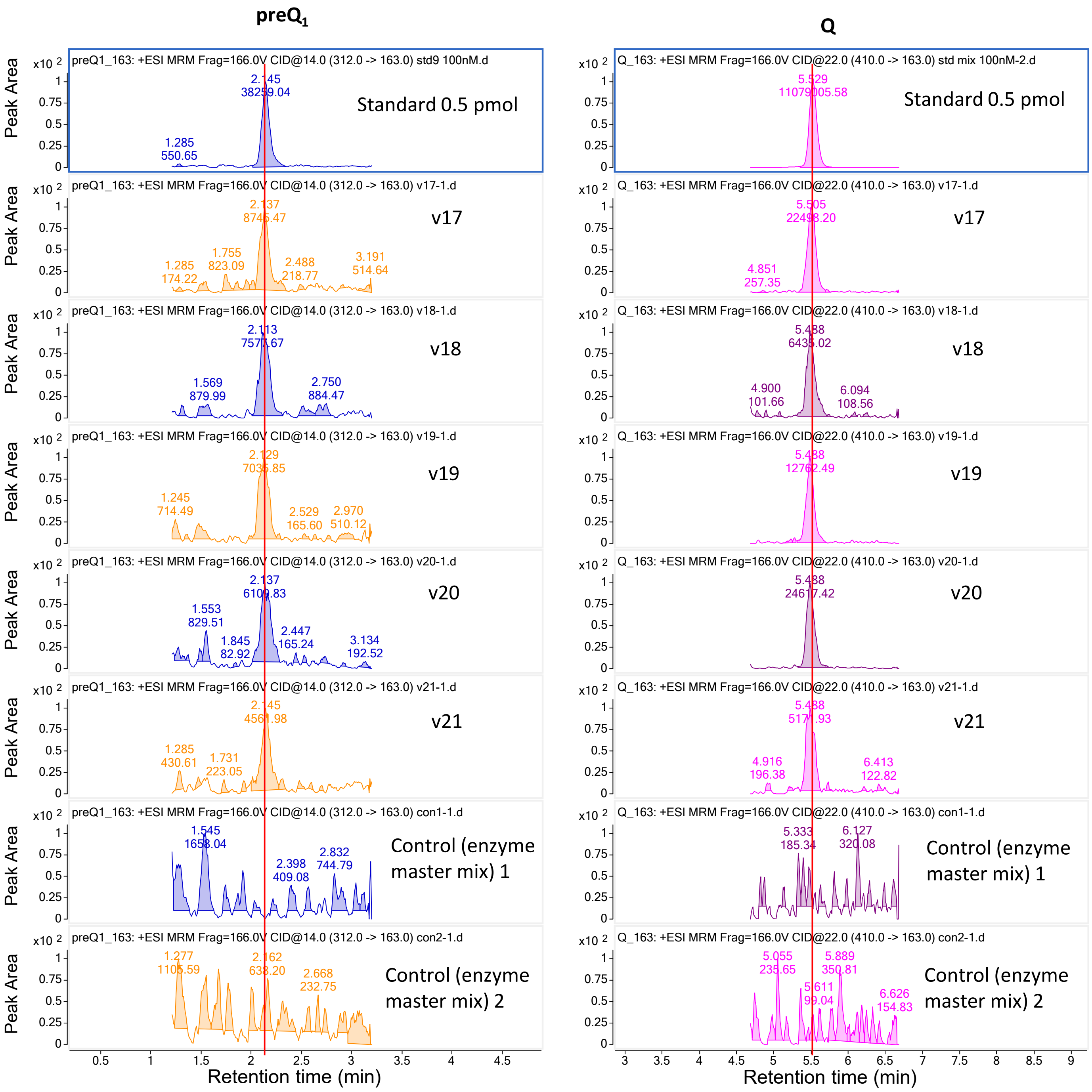


**Figure S3.** LC-MS analysis of tRNA modifications $preQ_1$ and Q in *B. henselae*. 600 ng of hydrolysate was injected. Enzyme master mix was used as an empty control. For each condition, two technical replicates were performed for each sample.

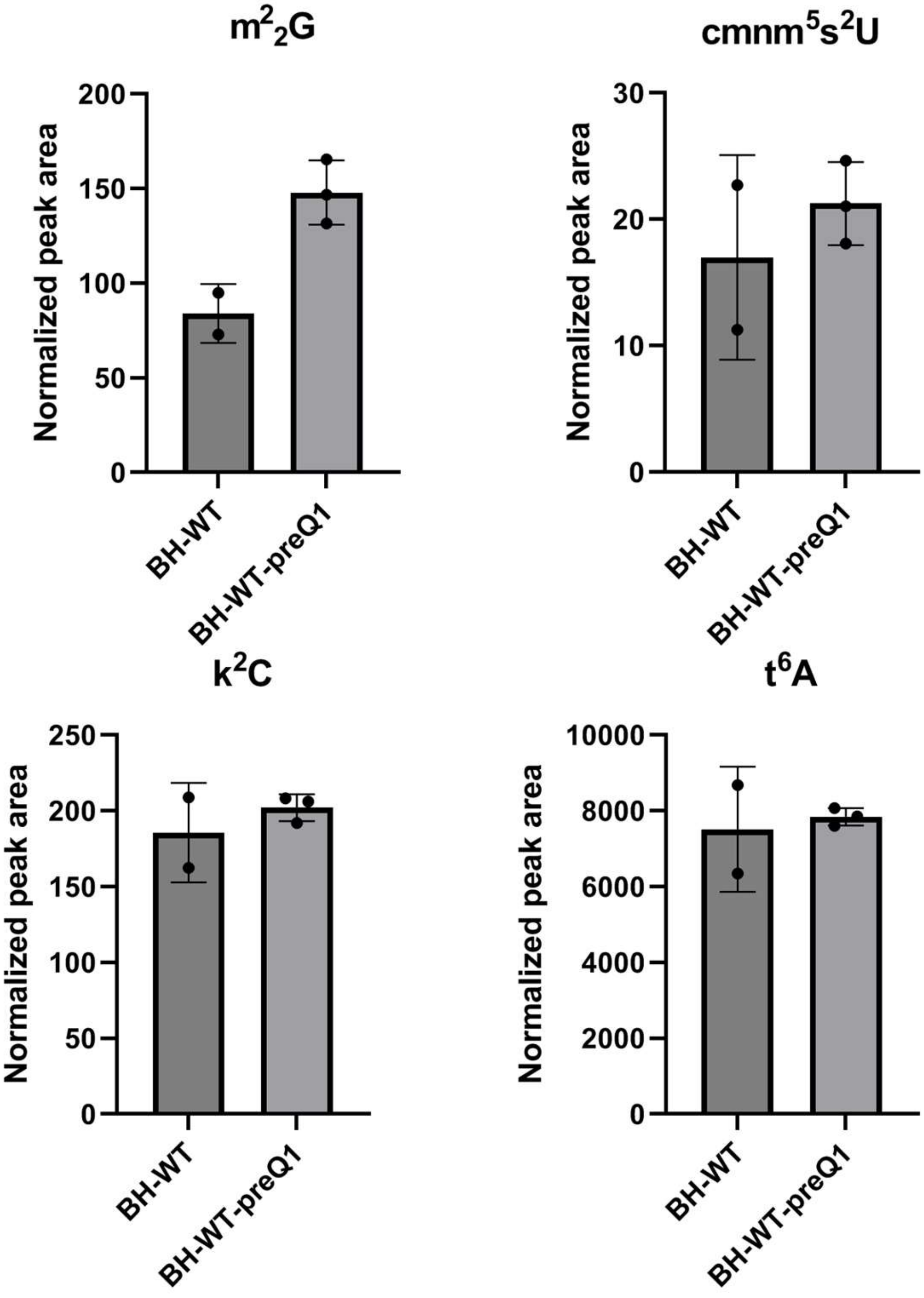


**Figure S4.** Quantification of $m^2_2G$, $cmnm^5s^2U$, $k^2C$ and $t^6A$ in *B. henselae*. LC-MS/MS analysis of modified ribonucleosides $m^2_2G$, $cmnm^5s^2U$, $k^2C$ and $t^6A$ was performed on 600 ng of hydrolyzed small RNA, with signal intensities normalized to the sum of UV absorbances of the canonical ribonucleosides, using an inline UV detector, to correct for differences in amounts of injected RNA. Data represent mean ± SD for three biological replicates of *B. henselae* BH-WT+preQ1 (BH-WT+preQ1) and mean ± deviation about the mean for two biological replicates of *B. henselae* BH-WT (BH-WT); technical duplicates were performed for each biological replicate. All modifications were validated with standards except $k^2C$, for which identification was based on retention time and MS-MS fragmentation.

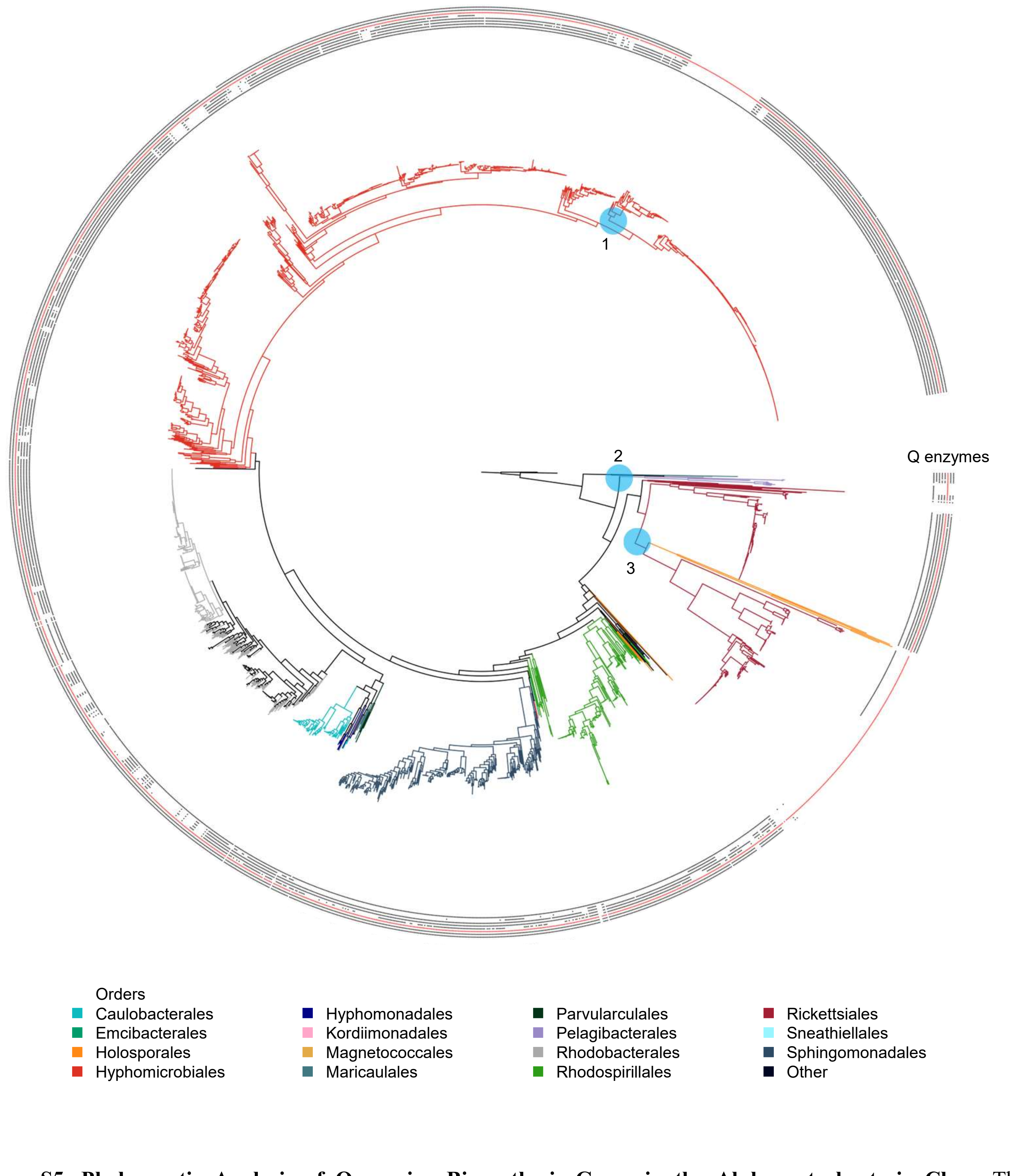


**Figure S5. Phylogenetic Analysis of Queuosine Biosynthesis Genes in the Alphaproteobacteria Class.** The maximum likelihood tree of 2,127 strains within the Alphaproteobacteria class. *E. coli* strains were used as the outgroup. Bootstrap values >0.5 are shown. The branches are colored by orders. The presence of Queuosine biosynthesis proteins is indicated by the circles: from inner to outer layers, FolE1/2, QueD, QueE, QueC, QueF, TGT (red), QueA, and QueG/H. The clades where the loss of Q pathway may occur are indicated in blue. #1 clade is presented in Fig. 6. #2 and #3 clades are presented in Fig. S7.

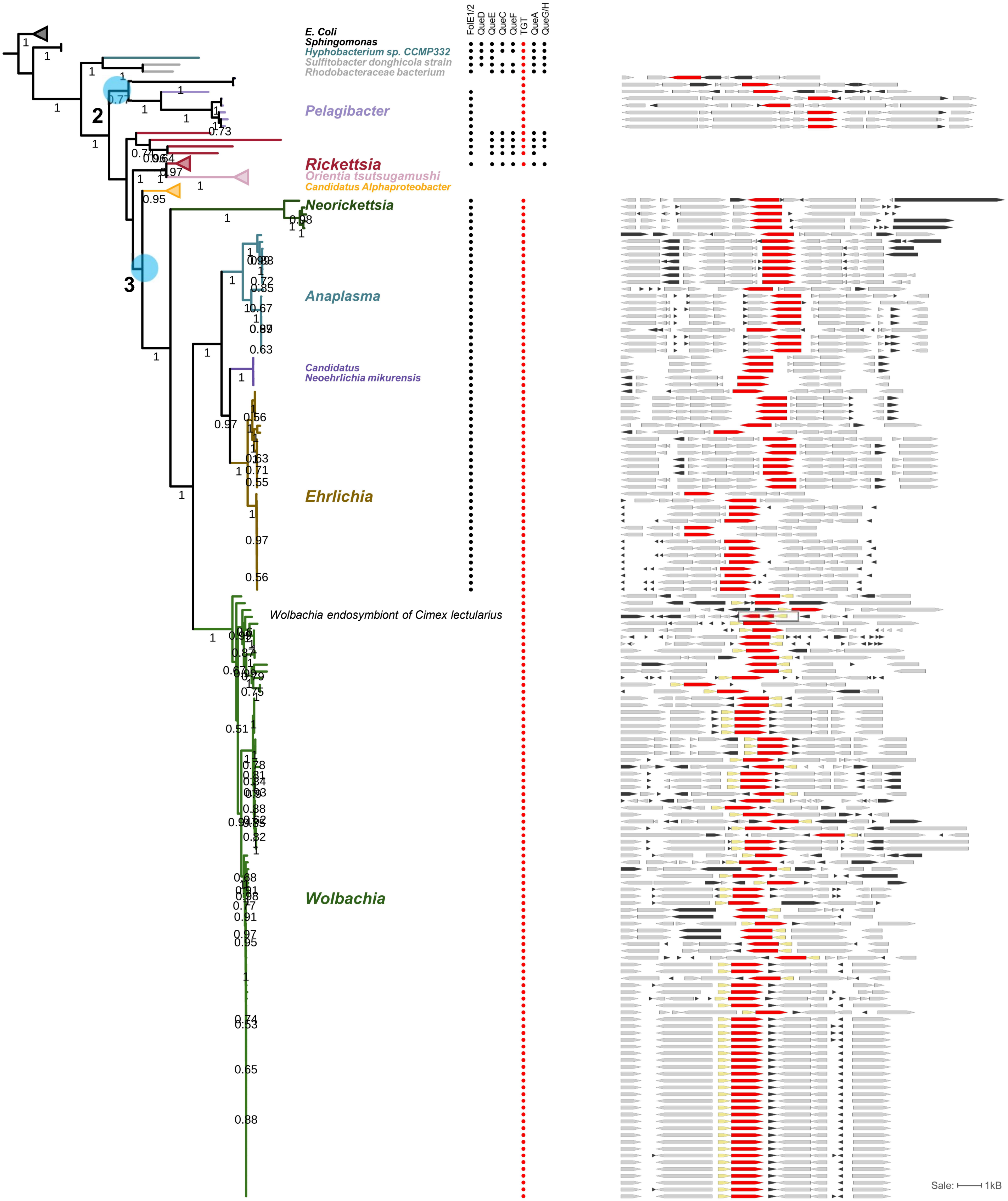


**Figure S6. Phylogenetic Analysis of Queuosine Biosynthesis Genes in the Rickettsiales order.** The clade of the *Rickettsiales* order from the tree of Alphaproteobacteria (Fig. 6A). The branches of *E. coli, Rickettsia*, *Orientia tsutsugamushi*, and *Candidatus Alphaproteobacter* were collapsed. The nodes where the loss of Q pathway may occur are indicated in blue. The presence of Queuosine biosynthesis proteins is indicated by the circles with TGTs highlighted in red. A comparative view of the corresponding truncated and full-length *tgt* gene variants in different strains in the tree is shown on the right. Red, *tgt*; yellow, NADH-ubiquinone oxidoreductase chain E (EC 1.6.5.3) encoding gene; black, hypothetical genes; gray, other genes. Gene IDs are provided in supplementary Table S4.
